# AnnotateAnyCell: Open-Source AI Framework for Efficient Annotation in Digital Pathology

**DOI:** 10.1101/2025.11.02.686114

**Authors:** Shourya Verma, Aditya Malusare, Mengbo Wang, Luopin Wang, Arpan Mahapatra, Abigail English, Abigail Cox, Meaghan Broman, Simone De Brot, Grant N. Burcham, Deborah Knapp, Deepika Dhawan, Mario Sola, Vaneet Aggarwal, Ananth Grama, Nadia Atallah Lanman

**Affiliations:** Computer Science, Purdue University; Industrial Engineering, Purdue University; Comparative Pathobiology, Purdue University; Veterinary Clinical Sciences, Purdue University; Purdue Institute for Cancer Research, Purdue University; Veterinary Pathology, University of Bern

**Author notes:** Equal Contribution (Shourya Verma, Aditya Malusare).

**Keywords:** Digital Pathology, Cell Annotation, Active Learning

## Abstract

Manual annotation of histopathological whole slide images remains a critical bottleneck for computational pathology and clinical AI deployment, requiring prohibitive expert time at scale. Here we present an open-source semi-supervised framework combining active contrastive learning with iterative human-in-the-loop feedback for efficient cellular annotation and classification. The pipeline integrates Cellpose segmentation, UMAP-based latent space visualization, and contrastive learning with pseudolabel propagation, evaluated on five whole slide images of canine invasive urothelial carcinoma across low, intermediate, and high histological grades at 40× magnification. Latent space clustering-guided annotation required 47 minutes compared to 63 minutes for sequential annotation, a 25% reduction (95% CI 18–32%). Classification accuracy reached 96.3% ± 1.2% for mitotic figures and 98.3% ± 1.4% for nucleoli using 1,075 labeled samples, with nucleoli classification achieving 95.5% ± 1.5% accuracy from only 215 samples. Inter-annotator agreement was high for chromatin (*κ* = 1.00) and nucleoli (*κ* = 0.95) but moderate for mitotic figures (*κ* = 0.58) and nuclear shape (*κ* = 0.36), reflecting intrinsic morphological ambiguity in these categories. This framework substantially reduces annotation burden while achieving expert-level accuracy for well-defined morphological features, providing a scalable path toward AI-assisted diagnostics in resource-constrained pathology settings.

## 1 Introduction

The integration of AI into digital pathology presents opportunities for systematic analysis of histopathological images at unprecedented scale [12, 29]. Modern whole slide images at 40× magnification contain hundreds of thousands of cells, typically taking several gigabytes of storage, and often exceeding the capacity for comprehensive manual review [28, 20]. While deep learning methods have shown promise for automated cellular analysis, their clinical deployment is limited by the requirement for extensive manual annotations by expert pathologists, who must spend hundreds of hours delineating cellular boundaries and assigning morphological classifications. Current approaches primarily use fully supervised learning, which requires extensive labeling of training datasets. This annotation burden is exacerbated by poor model generalization [25] across institutions due to variations in tissue preparation protocols and scanner characteristics, necessitating institution-specific retraining. Semi-supervised learning methods can leverage abundant unlabeled data alongside limited expert annotations. However, existing platforms require substantial computational infrastructure, lack intuitive interfaces for pathologist workflows, and do not provide iterative feedback loops to build clinical trust [1, 9, 19]. We developed AnnotateAnyCell, an open-source semi-supervised framework combining deep learning with an interactive web-based interface designed for pathologist workflows. The system enables annotation within learned embedding spaces, where morphologically similar cells naturally cluster, integrating active learning to intelligently select informative samples for expert review. We evaluated this framework on canine invasive urothelial carcinoma specimens, a translational model for human muscle-invasive bladder cancer [11], to determine annotation efficiency and classification accuracy for diagnostically relevant nuclear morphology features.

## 2 Background

### Cell Annotation

Desktop applications like QuPath [2] and Orbit [23] provide efficient tile-based visualization with manual annotation capabilities, while commercial platforms including HALO (Indica Labs) and Visiopharm dominate clinical deployment with FDA-cleared algorithms, validated workflows, and enterprise support infrastructure. Web-based platforms such as Cytomine [13] and Digital Slide Archive [8] enable collaborative annotation through RESTful APIs and distributed architectures, though they require substantial server infrastructure. Recent AI-assisted tools have dramatically improved annotation efficiency [27]: Quick Annotator [16] demonstrated significant speedups over manual annotation by training U-Net models concurrently during annotation and using UMAP-based patch selection for active learning, while PatchSorter [26] achieved improvements through deep learning-based object labeling [3].

### Active Learning

Active learning has emerged as a critical approach to address the substantial annotation burden in digital pathology. Recent work has demonstrated significant efficiency gains through intelligent sample selection strategies. Diversity-Aware Data Acquisition combined uncertainty and diversity clustering to achieve comparable AUC with 96% fewer patches for tumor-infiltrating lymphocyte classification [15]. Human-in-the-loop systems like Quick Annotator [16] have reduced annotation time through iterative review-and-revise workflows, where models provide hypothesized annotations that pathologists can rapidly verify. Uncertainty quantification methods, particularly Monte Carlo dropout [7], have enabled high-confidence predictions robust to domain shift. Despite these advances, most active learning systems operate at the patch or region level rather than individual cells, and few leverage modern embedding spaces for direct interactive annotation. AnnotateAnyCell significantly extends these approaches by combining active learning with contrastive learning-based embeddings in UMAP [14] space, enabling pathologists to annotate cells interactively across learned feature representations rather than spatial coordinates alone.

## 3 Methods

Our framework implements a semi-supervised active learning pipeline (Figure 1) for efficient cellular structure annotation through four iterative stages. First, deep learning models segment H&E-stained tissue images to extract individual cells as multi-modal representations: raw patches, isolated regions, and semantic masks. Second, extracted cells are presented to pathologists in a learned embedding space where morphologically similar cells cluster for efficient phenotype labeling. Third, a contrastive learning framework integrates expert labels and unlabeled data to generate high-confidence pseudolabels and prioritize informative samples for review. Fourth, a multi-modal autoencoder processes labeled and pseudolabeled data to learn compact latent embeddings capturing fine-grained morphology and higher-level context, producing updated visualizations. This loop iterates until sufficient annotation coverage is achieved, progressively improving model accuracy and expert efficiency.

**Figure 1:**
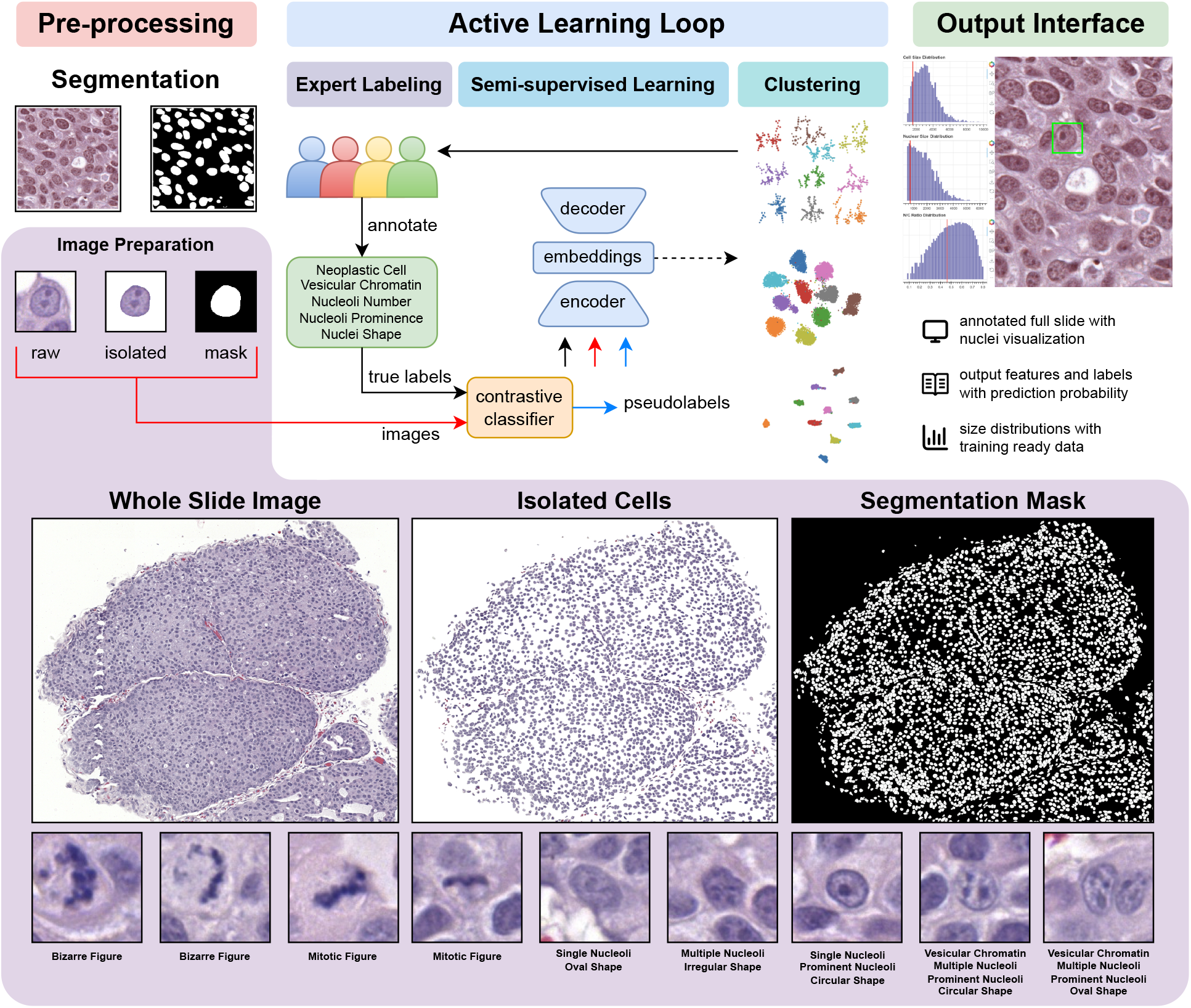
**Pre-processing stage:** Raw H&E-stained tissue images undergo segmentation using Cellpose models to generate nuclear masks. Each detected nucleus is extracted as three complementary representations: raw image patches, isolated nuclear regions, and semantic masks; **Active Learning Loop:** Expert pathologists (represented by colored figures) provide initial annotations for nuclear morphology features of neoplastic cells, including mitotic figures, vesicular chromatin, and prominent nucleoli. These labeled samples train a contrastive classifier that generates embeddings and pseudolabels for unlabeled data. A convolutional autoencoder processes the multi-modal inputs (raw, isolated, mask) to learn compact latent representations; **Output Interface:** The system produces annotated whole slide images with nuclei visualization, quantitative feature distributions, and prediction probabilities for downstream analysis. The iterative process refines the model’s understanding of cellular morphologies through expert feedback. **Image Preparation:** Left: Representative whole-slide image (WSI). Center: Nuclear segmentation results using pretrained Cellpose models. Right: Binary segmentation mask showing all detected valid cells. (Bottom row) Nine examples of 128×128 pixel cellular tiles illustrating morphological diversity across expert-defined annotation categories. Examples include bizarre mitotic figures showing irregular mitotic configurations, canonical mitotic figures demonstrating typical mitotic chromatin condensation patterns, nucleolar variations (single vs. multiple nucleoli) combined with nuclear shape classifications (circular, oval, irregular), and chromatin texture patterns (vesicular chromatin).

### Dataset and Sample Collection

We evaluated our framework on canine invasive urothelial carcinoma (IncUC), a spontaneous tumor model [11, 6] that recapitulates human muscle-invasive bladder cancer biology for translational research. Tissue samples from client-owned dogs with naturally occurring IncUC were collected at Purdue University Veterinary Teaching Hospital under informed consent and institutional protocols. Formalin-fixed paraffin-embedded blocks were sectioned at 4 *µm* and H&E-stained using automated protocols (Leica Autostainer XL). Slides were digitized at 40× magnification (0.25 *µm/*pixel, Aperio AT2), producing whole slide images of 80,000×100,000 pixels. Cases were selected for diagnostic quality (minimal artifacts, adequate fixation, uniform staining) and tumor cellularity (>30% neoplastic nuclei). Five representative slides from necropsy samples captured morphological diversity with mitotic activity ranging from 5 to >30 mitoses per 10 high-power fields and papillary or solid architectural patterns. ACVP Board-certified veterinary pathologists defined annotation categories based on diagnostically relevant nuclear features: mitotic figures (canonical and atypical configurations), nucleolar characteristics (absent, single, multiple, prominent), chromatin texture (vesicular vs. hyperchromatic), and nuclear shape (circular, oval, irregular). These features were selected for their prognostic significance in canine and human urothelial carcinoma [17, 18], relevance to grade assignment, and reliable inter-observer agreement.

### User Interface

The interactive interface was built with BokehJS (JavaScript with Python bindings) for real-time multi-user annotation with persistent sessions and checkpoint functionality. The primary workspace (Fig. 2) comprises three coordinated panels: the central panel displays a UMAP [14] embedding as an interactive scatter plot where each point represents a cellular tile, color-coded by status (blue: unlabeled, green: labeled, orange: selected), with hovering revealing original tiles, segmented regions, and WSI coordinates; the left panel provides stepwise protocol guidance with interactive analytics including histograms of nuclear area and nuclear-to-cytoplasmic ratios that dynamically update with selections; the right panel presents 512×512 high-resolution previews with annotation controls for pathologist-defined morphological classes (mitotic figures, chromatin patterns, nucleoli, nuclear shape), recording confidence scores and timestamps for inter-observer variability analysis. Real-time progress tracking displays dataset coverage, and the “Update Clusters” button triggers asynchronous model retraining, automatically updating the embedding to reveal refined clusters. The results interface (Fig. 3) reuses the three-panel layout with annotation count summaries, WSI display with cyan markers for annotated cells supporting pan/zoom exploration, and detailed individual annotation inspection. Data export via CSV and advanced filters enable hypothesis generation. Eleven board-certified veterinary pathologists tested the interface by labeling 300 cells each to evaluate usability and inter-annotator agreement.

**Figure 2:**
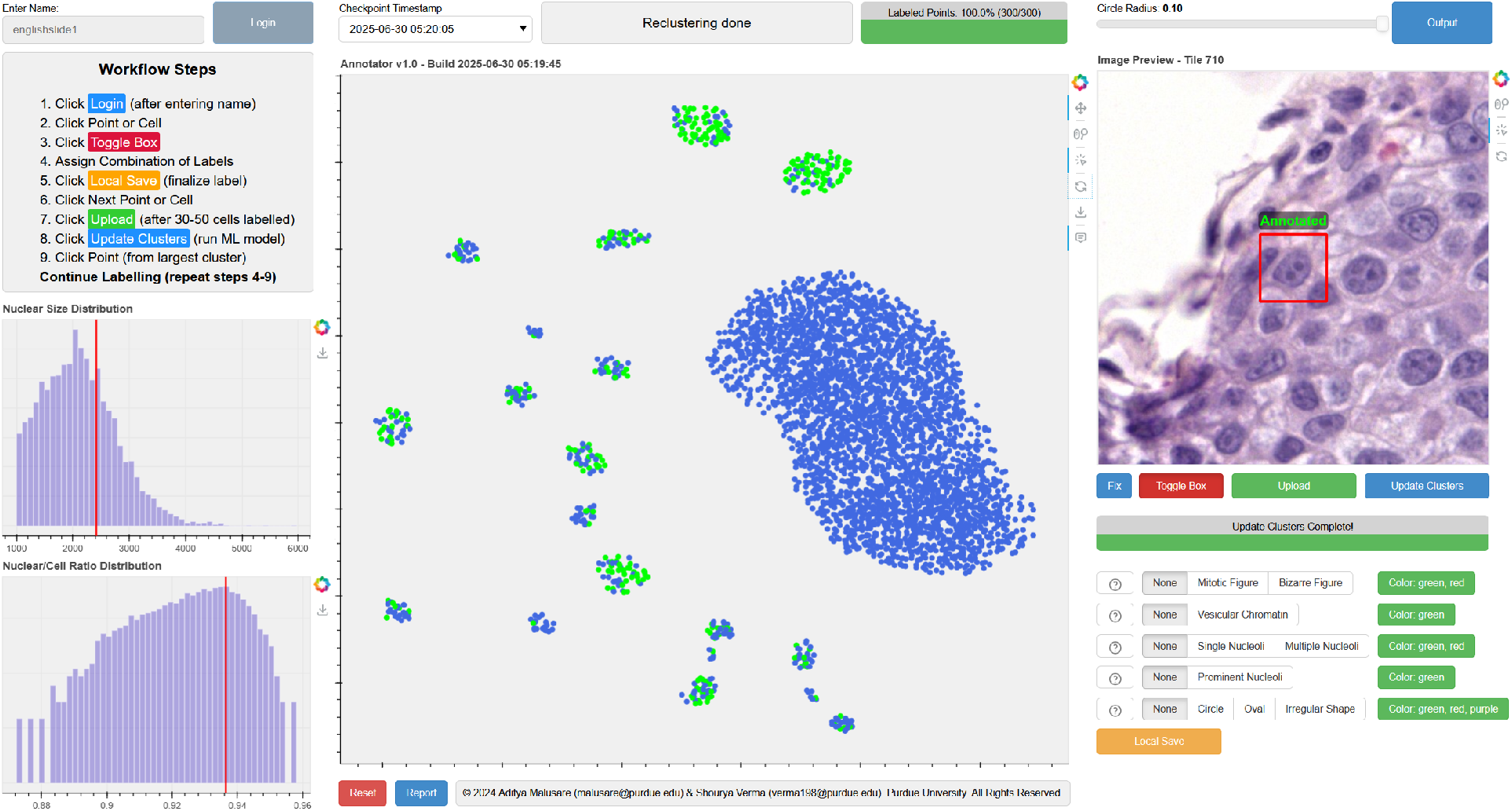
Left panel: Displays workflow guide for pathologists, nuclear size distribution histogram, and nuclear-to-cell ratio distribution with red indicators showing the currently selected cell’s measurements. Center panel: Interactive UMAP embedding visualization, where each point represents a cellular tile extracted from the whole slide image. Cells are color-coded based on annotation status (blue: unlabeled, green: labeled, orange: selected) and naturally cluster based on morphological similarities. Right panel: Shows a high-resolution preview of the selected cell tile (Tile 710) with morphological annotation options including mitotic figures, chromatin patterns, nucleoli characteristics, and nuclear shape classifications. The interface includes progress tracking (300/300 labeled points), version control, and model retraining capabilities (“Update Clusters” button).

**Figure 3:**
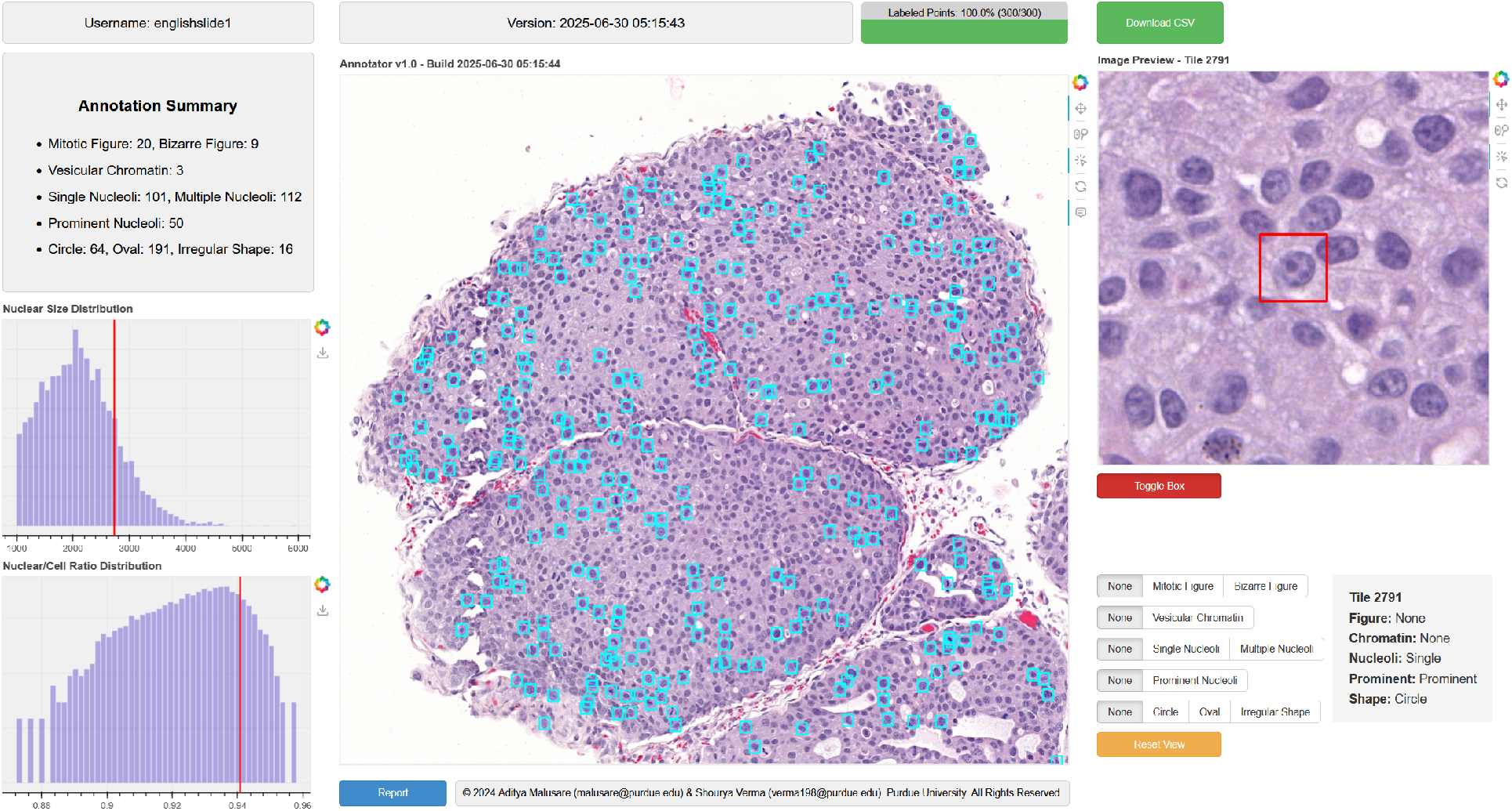
Left panel: Displays annotation summary statistics including counts for different morphological features along with distribution histograms. Center panel: Whole slide image view with cyan markers indicating the spatial locations of all annotated cells across the tissue sample, enabling pathologists to assess spatial distribution patterns and tissue architecture context. Right panel: Individual cell inspection tool showing detailed annotations for the selected cell (Tile 2791) with its specific morphological classifications. The interface includes data export functionality (Download CSV), version information, and the ability to filter and visualize cells based on specific morphological criteria. This output interface enables analysis of annotation results and supports research workflows requiring quantitative cellular phenotype data.

### Image Pre-processing

We use pre-trained Cellpose models [22]: cyto3 [21] for cell and nuclear segmentation. Segmentation masks are filtered to remove artifacts, partial objects, and merged cells. Cell and nuclear masks are integrated, enforcing containment (each nucleus within a cell body); cells without detected nuclei are treated entirely as nuclei. From segmented regions, we extract tile patches: 128×128 pixels for training and 512×512 pixels for expert inspection. Three versions per tile are generated: (i) raw WSI image, (ii) semantic mask, and (iii) isolated cell/nucleus with surrounding tissue removed, all bilinearly interpolated to standard dimensions. Metadata including nuclear-to-cytoplasmic ratio, nuclear and cellular areas, and centroid coordinates are stored for analysis and annotation context. UMAP [14] is applied to 128×128 patches for dimensionality reduction, enabling visualization and clustering of morphologically similar cells.

### Active Learning Loop

To minimize annotation burden, we developed a semi-supervised active learning pipeline iteratively selecting informative samples for expert review. The process begins with UMAP embeddings of 128×128 cellular tiles creating a 2D morphological feature space for pathologists to explore and annotate representative tiles, stored in a MongoDB database with secure access, version control, and multi-expert collaboration support. A contrastive learning model with convolutional backbone, projection head, and classification head is trained, combining supervised classification loss on labeled data with unsupervised contrastive loss on all samples to embed similar morphologies together while separating dissimilar ones. The trained model generates class-balanced pseudolabels by selecting top-N highest-confidence samples per class to avoid bias toward dominant morphologies. For subsequent rounds, uncertainty sampling with diversity promotion identifies informative examples. A convolutional autoencoder trained on annotated and pseudolabeled data processes raw images, nuclear features, and semantic masks simultaneously, capturing fine-grained morphology and contextual structure in its latent space through skip connections and a variational bottleneck. UMAP projection of these embeddings yields updated visualizations clustering morphologically similar cells while preserving expert-defined boundaries. Experts use this refined space to validate or correct pseudolabels and annotate uncertain regions, iteratively refining the nuclear morphology space and improving classification accuracy, annotation efficiency, and novel phenotype discovery.

### Pseudolabel Generation

Our semi-supervised learning framework integrates supervised and unsupervised techniques through three components: a convolutional classifier for feature extraction, a contrastive learning module for representation enhancement, and a selective pseudolabeling strategy for label propagation. The base CNN extracts features using four convolutional blocks (Conv2D with 3 × 3 kernels, ReLU, max-pooling), increasing channel depth (16 → 128) while reducing spatial dimensions, followed by four fully connected layers (1024 → 128 neurons) with batch normalization and 50% dropout, with the final layer mapping to expert-defined nuclear morphology classes. A projection head maps CNN features to a 128-dimensional normalized embedding space for contrastive learning. The supervised classification objective combines with InfoNCE loss [4]:

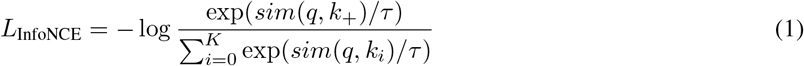

where *q* is the query embedding, *k*_+_ the positive key, *k*_*i*_ negative keys, *sim*(*u, v*) cosine similarity, *τ* temperature, and *K* number of negatives. The total objective is:

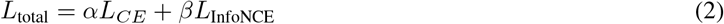

with *α* and *β* controlling loss balance. For pseudolabeling, the trained model uses class-balanced selection, choosing top-*N* most confident predictions per class (e.g., *N* = 20) to ensure balanced representation and mitigate bias toward dominant classes, with unselected samples retaining null labels and contributing only to unsupervised learning. Experimental validation [24] shows this approach outperforms fully supervised baselines and naive pseudolabeling.

### Autoencoder Model

We implement a multi-modal convolutional autoencoder with three input channels: raw cellular images, nuclear regions, and semantic masks. Each modality passes through parallel encoders (four Conv2D layers: 3 × 16 → 16 × 32 → 32 × 64 → 64 × 128, ReLU) with skip connections preserving spatial detail. Encoded features are flattened, passed through fully connected layers (1024 → 512, batch normalization), concatenated with one-hot pseudolabels, and processed through additional layers (1024 → 512). Following the VAE framework [10], mean and logvar projections map to a 256-dimensional latent space with reparameterization trick applied during training. Decoders mirror encoder structure with transposed convolutions. The training objective combines reconstruction and feature prediction losses:

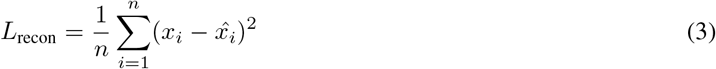

where *x*_*i*_ is input and 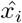 reconstruction, and feature prediction uses cross-entropy:

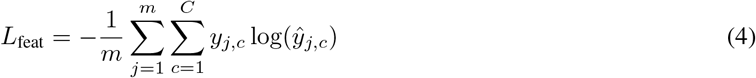

The total loss is:

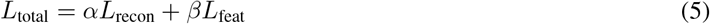

with *α, β* controlling trade-offs. The encoder transforms 128 × 128 × 9 input tensors into 256-dimensional embeddings capturing supervised annotation signals and unsupervised data structure, projected to 2D via UMAP [14] for interpretable visualization, improving representation quality, reconstruction fidelity, and clustering performance.

## 4 Results

### Annotator Cohort

The annotator cohort consisted of ACVP (American College of Veterinary Pathologists) and FMH (Foederatio Medicorum Helveticorum) board certified pathologists with more than 10 years of experience each. Annotations were also provided by the pathology residents in training, which were verified by the certified pathologists.

### Model Performance

In Figure 4 **A**. unsupervised UMAP projection of 128×128 pixel cellular tiles creates a 2D morphological landscape. At initialization (0 points), all cells are unlabeled. With 50 labeled points, annotations scatter sparsely as pathologists explore diversity and establish class boundaries. At 100-200 points, annotation density increases in coherent neighborhoods via uncertainty sampling with diversity promotion. At 3000 points, labeled cells (green) populate discrete clusters and boundaries, with a large central cluster of unlabeled cells not yet confidently assigned and peripheral clusters capturing annotated dominant and rare phenotypes. The framework aims to break the large unlabeled cluster into phenotypically necessary sub-clusters. Bottom row shows feature-specific visualization of the 300-point embedding colored by morphological classifications, revealing feature-dependent clustering quality.

**Figure 4:**
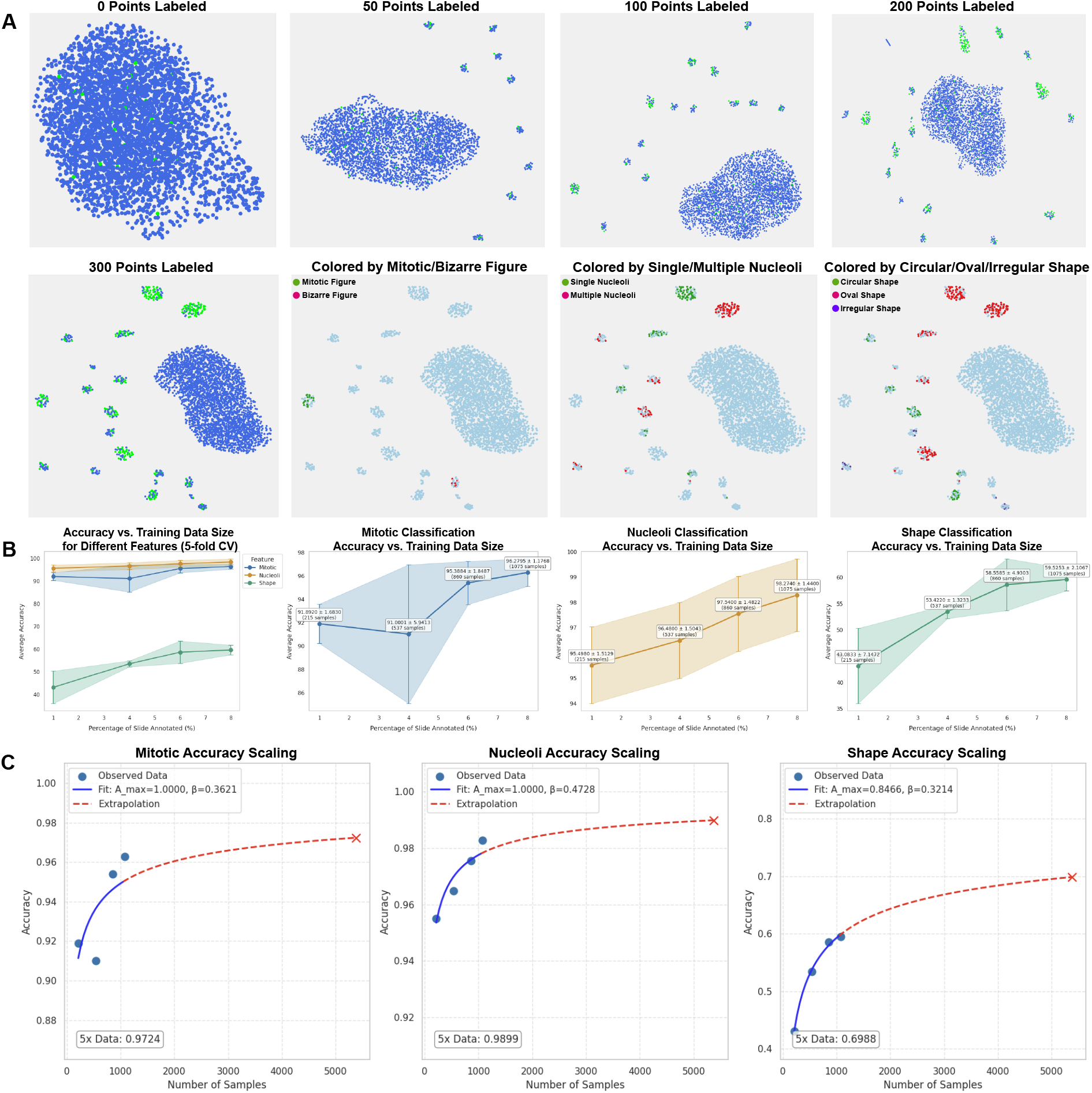
**A.** Sequential snapshots of the learned embedding space progression of annotations (green) within the unlabeled cellular population (blue). **B**. Classification evaluation of mitotic figures, nucleoli, and nuclear shape across training sizes using 5-fold cross-validation. **C**. Scaling behaviors highlight fundamental differences in data efficiency.

Figure 4 **B**. shows nucleoli classification achieved highest accuracy (98.3% at 1075 samples), mitotic reached 96.3%, and shape plateaued at 59.5%. Mitotic classification showed strong baseline (91.9% ± 1.7% at 215 samples), transient dip at 4% coverage (91.0% ± 5.9%), recovery to 95.4% ± 1.8% at 6%, and plateau at 96.3% ± 1.2% with 1075 samples. Mid-range instability likely reflects class imbalance or sampling variability. Above 800 samples, performance stabilized with narrow bounds, exceeding the 64% inter-annotator agreement. Nucleoli classification started at 95.5% ± 1.5% with 215 samples and improved monotonically to 98.3% ± 1.4%. Consistently narrow confidence intervals mirror the 95% inter-annotator agreement, with final accuracy approaching the theoretical ceiling. Shape classification proved most difficult, rising from 43.1% ± 7.1% to 59.5% ± 2.1% at full data. Despite the largest absolute gain (16.4 points), performance remained lowest due to subjectivity of categorical shape labels and dependence on continuous morphological spectra with complex contextual factors. Figure 4 **C**. Shows power-law fits highlight distinct scaling regimes: nucleoli and mitotic figures exhibit early saturation with diminishing returns, whereas shape maintains a steep learning trajectory, suggesting substantially more labeled data would be required to approach parity.

### User Performance

Figure 5 summarizes spatial agreement maps, temporal efficiency, feature-level metrics, annotator reliability, inter-feature correlations, and disagreement structures. In Figure 5 **A**. clustering-based active learning substantially outperformed sequential annotation, requiring 47 vs. 63 minutes for 300 cells (25% reduction; 16 minutes absolute; 9.4 vs. 12.6 sec/cell) averaged over 11 users, with consistent improvements across volumes: 22% at 50 cells (18 vs. 23 min), 30% at 100 cells (26 vs. 37 min), and 25% at 150 cells (33 vs. 44 min), all scaling linearly without fatigue effects. Efficiency gains stem from embedding-based grouping of 5-10 morphologically similar cells, reducing cognitive load and context switching, with interactive UMAP visualization, real-time statistics, and uncertainty- and diversity-based sampling ensuring informative morphologies are prioritized. Spatial annotation patterns revealed feature-specific reliability: mitotic figures were sparse (38 sites: 25 concordant, 13 disagreements at 0.2-1.0 intensity), reflecting difficulty distinguishing mitotic events from apoptotic bodies; nucleoli achieved highest density with uniform coverage and minimal disagreements (16 sites), mostly borderline single vs. multiple classifications; nuclear shape showed highest absolute disagreement with moderate disagreements (0.4-0.6) clustered regionally and sporadic high-intensity disagreements (≥ 0.8) reflecting divergent classification strategies, with no regional clustering indicating cell-level ambiguity rather than slide-level artifacts.

**Figure 5:**
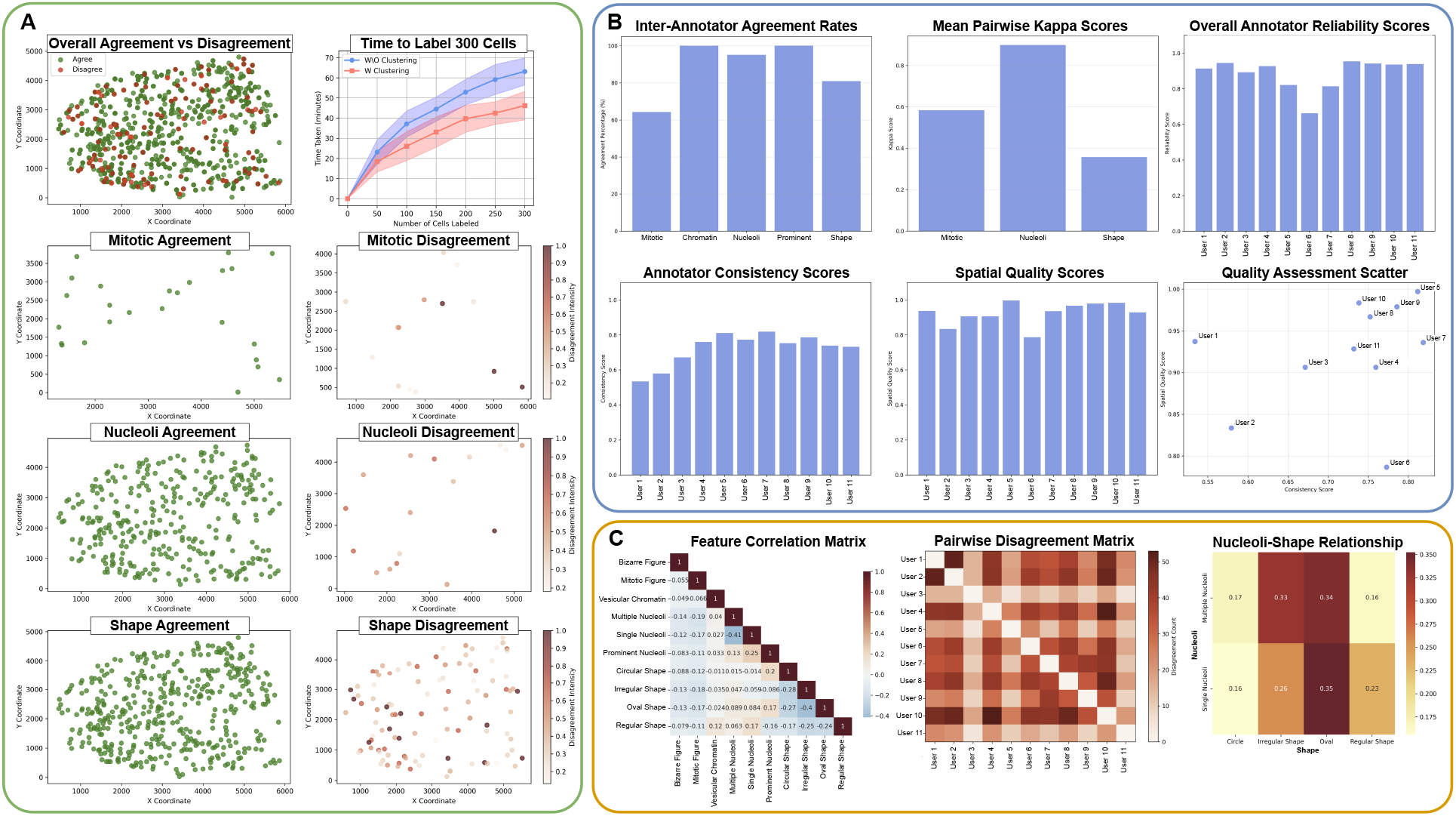
**A.** Illustrates labelling time with and without clustering, along with spatial distributions of agreement (green) and disagreement (brown) reveal localized cell-level ambiguities, with disagreement intensity encoded by saturation. **B**. Presents the annotation workflow metrics like agreement rates, reliability scores, consistency and spatial coverage scores across 11 annotators. **C**. Illustrates feature correlation demonstrating nuclear features are largely independent dimensions of morphology, and pairwise disagreement analyses showing some users might disagree more than others.

Figure 5 **B**. shows agreement rates varied substantially: chromatin texture and prominent nucleoli achieved perfect concordance (100%), nucleoli classification showed high agreement (95%, Cohen’s *κ* = 0.95) [5], shape was variable (81%, *κ* = 0.36), and mitotic activity most challenging (64%, *κ* = 0.58). Reliability across 11 annotators ranged from 0.65-0.95 (Users 2, 8, 9, 10 achieved ≥ 0.95; User 6 lowest at 0.65). Decision consistency ranged from 0.52-0.82 (User 5: 0.82; Users 1, 2: 0.52-0.57), and spatial quality scores were 0.80-1.00 (User 5: 1.00; User 6: 0.80). High performers (Users 5, 9, 10) combined strong consistency (0.75-0.82) with near-perfect spatial coverage (0.95-1.00), while Users 2 and 6 emerged as retraining candidates. Chromatin and nucleoli are robust for clinical and computational pipelines, while mitotic activity and shape require standardized protocols or noise-robust learning.

Figure 5 **C**. shows pairwise correlations were weak (|*r*| < 0.3), with strongest between single and prominent nucleoli (*r* = 0.25), oval shape weakly correlated with single (*r* = 0.084) and prominent nucleoli (*r* = 0.17), irregular shape negatively correlated with circular (*r* = − 0.27) and oval (*r* = − 0.40), and multiple and single nucleoli mutually exclusive (*r* = − 0.41). Cross-tab analysis revealed biologically meaningful associations: multiple nucleoli enriched in oval (0.34) and irregular (0.33) nuclei; single nucleoli frequent in oval (0.35) and irregular (0.26), with both nucleolar states elevated in irregular nuclei, suggesting nuclear irregularity as an independent atypia marker supporting a multidimensional diagnostic model. Inter-annotator disagreement revealed systematic variability: Users 1, 4, 10 showed highest discord (>40 with multiple annotators); Users 3, 9 showed strong concordance; asymmetric patterns revealed two annotation “schools”: concordant (Users 2, 3, 6, 9) and discordant (Users 1, 4, 10). For machine learning, features with near-perfect agreement (chromatin, nucleoli) provide robust ground-truth, while ambiguous features (mitotic figures, shape) require consensus labeling or noise-tolerant algorithms, with disagreement intensity as spatial uncertainty enabling principled annotation weighting.

## 5 Discussion

We presented a semi-supervised active learning framework for efficient nuclear morphology annotation in histopathological images, integrating contrastive learning, class-balanced pseudolabeling, and multi-modal variational autoencoders. Applied to canine invasive urothelial carcinoma, the framework achieved 25% reduction in annotation time (47 vs. 63 minutes for 300 cells) through clustering-based sample selection and interactive UMAP visualization. Classification performance reached 98.3% for nucleoli, 96.3% for mitotic figures, and 59.5% for nuclear shape, with nucleoli achieving 95.5% accuracy using only 215 labeled samples. The InfoNCE loss successfully leverages unlabeled data to establish morphological feature spaces where similar cells cluster, enabling robust classification from sparse annotations. The class-balanced pseudolabel generation proved effective, with performance improving without biased pseudolabel degradation. The multi-modal autoencoder processing raw images, nuclear features, and semantic masks through a 128-dimensional normalized embedding space captures diverse morphological characteristics. Integration of labeled and pseudolabeled data through the variational bottleneck creates smooth latent representations supporting both supervised classification and unsupervised structure learning. The iterative active learning loop demonstrates steep learning curves for mitotic and shape features, with updated embeddings revealing refined morphological relationships guiding subsequent annotation rounds.

Inter-annotator analysis revealed feature-dependent reliability: chromatin texture and prominent nucleoli achieved perfect agreement (100%), while mitotic activity (64%, *κ* = 0.58) and nuclear shape (81%, *κ* = 0.36) showed higher variability. Weak pairwise feature correlations (|*r*| < 0.3) support a multidimensional diagnostic model where features contribute complementary information. For clinical deployment, differential performance suggests tiered confidence thresholds: nucleoli and mitotic classifications (96-98% accuracy) achieve reliability for automated screening or diagnostic support, while shape classifications (59% accuracy) require human verification or uncertainty quantification. Clinical workflows could automatically accept high-confidence predictions while flagging ambiguous cases for expert review. Data efficiency demonstrated by achieving 95.5% accuracy with 215 samples supports rapid institutional adaptation. Pathologists could annotate 200-500 representative local cells to generate institution-specific models capturing local preparation protocols and scanner characteristics while leveraging general morphological knowledge from unsupervised pretraining. This democratizes computational pathology for resource-constrained settings without prohibitive annotation infrastructure. Future work will address label noise tolerance for ambiguous morphologies, extend the framework to multi-tissue generalization, and validate clinical utility in prospective diagnostic workflows. This human-in-the-loop approach demonstrates that combining expert knowledge with semi-supervised learning achieves expert-level performance while substantially reducing annotation burden.

## 6 Author Contributions

S.V. and A.M. created the AnnotateAnyCell platform and wrote the manuscript text, and created the figures.

M.W., L.W., A.M. performed data preprocessing and dataset curation.

A.E., A.C., M.B., S.D., G.B., D.K., D.D., M.S., N.L. performed annotations on the dataset using the tool as users.

V.A., A.G. guided the project as senior supervisors.

All authors reviewed the manuscript. Nadia Atallah Lanman is the corresponding author.

## 7 Acknowledgement

## 11 Data Availability

The datasets and the code will be open sourced on **Github:** https://github.com/shouryaverma/AnnotateAnyCell and zenodo for academic use only. Link to data will be provided upon acceptance.

